# Musculoskeletal fat imaging and quantification by high-resolution metabolite cycling magnetic resonance spectroscopic imaging at 3 T: A fast method to generate separate distribution maps of lipid components

**DOI:** 10.1101/732016

**Authors:** Ahmad A. Alhulail, Debra A. Patterson, Pingyu Xia, Xiaopeng Zhou, Chen Lin, M. Albert Thomas, Ulrike Dydak, Uzay E. Emir

## Abstract

**Purpose:** To provide a rapid, non-invasive fat quantification technique capable of producing separate lipid component maps.

**Methods:** The calf muscles in 5 healthy adolescents (age 12-16 years; BMI = 20 ± 3 Kg/m^2^) were scanned by two different fat fraction (FF) quantification methods. A high-resolution, density-weighted concentric ring trajectory (DW-CRT) metabolite cycling (MC) magnetic resonance spectroscopic imaging (MRSI) technique was implemented to collect data with 0.25 mL resolution within 3 minutes and 16 seconds. For comparative purposes, the standard Dixon technique was performed. The two techniques were compared using structural similarity (SSIM) analysis. Additionally, the difference in the distribution of each lipid over the adolescent calf muscles was assessed based on the MRSI data.

**Results:** The proposed MRSI technique provided individual FF maps for eight musculoskeletal lipids identified by LCModel analysis (L09, L11, L13, L15, L21, L23, L53, and L55) with mean SSIM indices of 0.19, 0.04, 0.03, 0.50, 0.45, 0.04, 0.07, and 0.12, respectively compared to that of Dixon’s FF map. Further analysis of voxels with zero SSIM demonstrated an increased sensitivity of FF lipid maps from data acquired using this MRSI technique over the standard Dixon technique. The trend of lipid spatial distribution over calf muscles was consistent with previously published findings in adults.

**Conclusion:** The advantages of this MRSI technique make it a useful tool when individual lipid FF maps are desired within a short scanning time.

## 1 INTRODUCTION

The accumulation of adipose tissue in the human body is a risk factor for many common health disorders, such as type 2 diabetes mellitus (T2DM) and cardiovascular disease.^1,2^ Indeed, obese children and adolescents are more likely to develop such health problems.^3–5^ It has been found that intramyocellular triglycerides (lipids) content has a direct relationship to insulin resistance,^6,7^ and is hypothesized to be a precursor to T2DM.^8,9^ Thus, a non-invasive method to reliably evaluate particular lipid content can be useful in the early detection and prevention of such diseases in children, adolescents, and adults.

Currently, there are several methods available to investigate fat content within the muscles of the body. A biopsy specimen can be extracted from the region of interest (ROI) and quantified in vitro by electron microscopy,^10^ histochemistry (Oil red O staining),^11^ or biochemical analysis.^12^ However, this method is invasive and does not provide the spatial distribution of lipids. Alternately, computed tomography (CT)^13^ and magnetic resonance imaging (MRI) can noninvasively image a larger area for fat content assessment. However, MRI is preferred over CT for fat quantification, especially in children, because CT uses ionizing radiation.

The Dixon MRI technique has been used to quantify in vivo total fat fraction (FF).^14^ This technique has been considered to be the standard MRI method in providing information about the fat level from a relatively large area in the body. In particular, this technique has been used to image an entire cross-section of the leg to assess elevated total fat content levels, which can be associated with Duchenne muscular dystrophy,^15^ tendon tear severity,^16^ Charcot-Marie-Tooth type 2 F disease,^17^ osteoporosis,^18^ and knee osteoarthritis.^19^

The available Dixon MRI techniques are utilized to map the total fat fraction rather than the individual lipid component (Table 1) as these techniques cannot differentiate the signal amplitude of each fatty acid separately. In certain situations, individual lipid component information might reflect different pathology or physiology. For instance, the methylene lipid group, which consisted of the intramyocellular lipid that resonates at 1.3 ppm (L13) and the extramyocellular lipid that resonates at 1.5 ppm (L15), is considered to evaluate arterial stiffness.^25^ However, in other situations, only the L13 is the lipid of particular interest since its elevated level has been found to be a biomarker for insulin resistivity^6,26^ as well as for mitochondrial disorder MELAS.^27^ In addition, it may indicate physiological information as it has been shown that muscle exercises result in altering the levels of L13.^28^ Similarly, the increase of saturated lipids and decline of unsaturated olefinic lipid signal in bone marrow was used as a sign of osteoporosis.^29^ All these studies indicate a need to measure the lipid components.

**TABLE 1.**
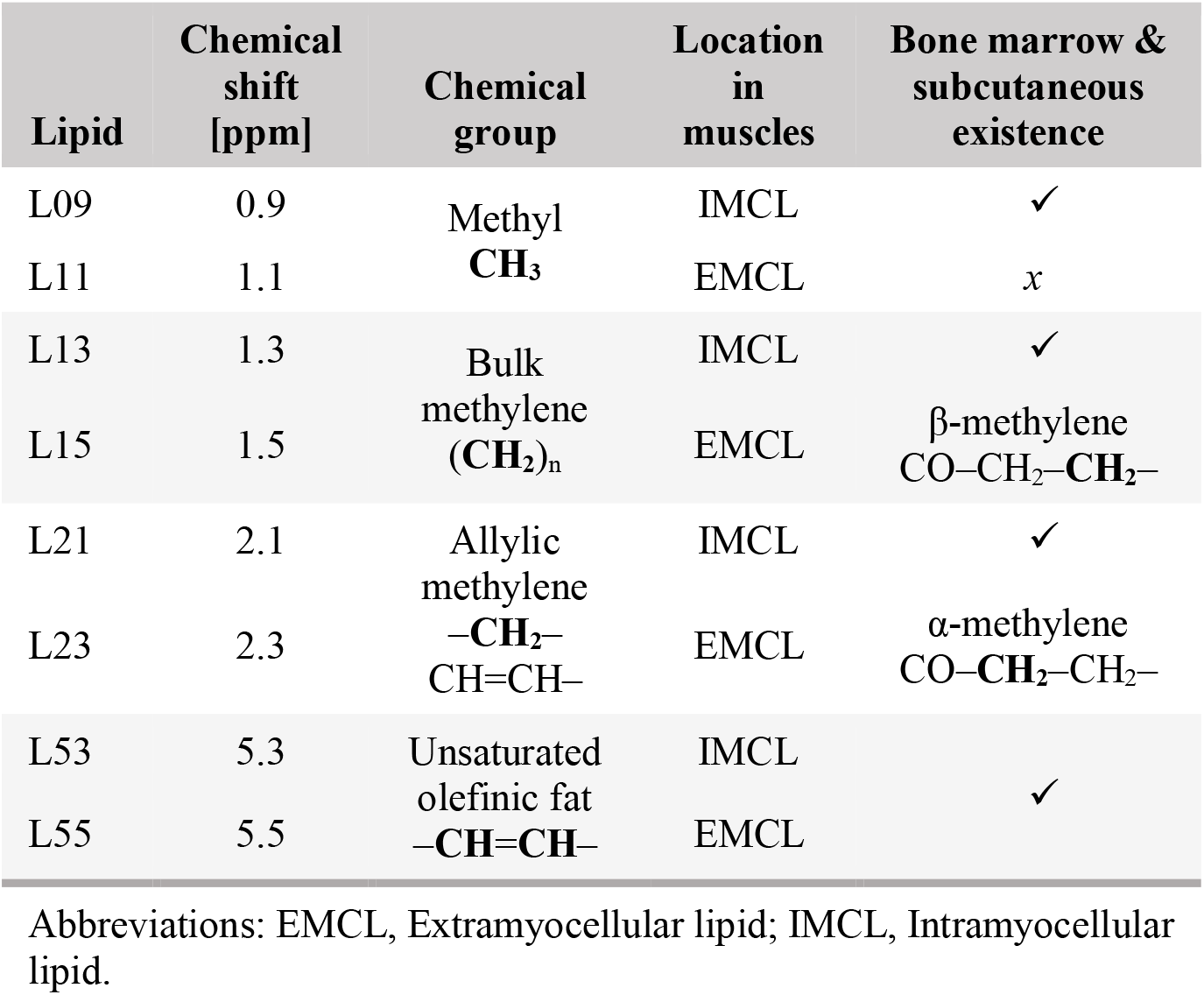
Common musculoskeletal lipid signals detected by MR techniques^20–24^

The single-voxel magnetic resonance spectroscopy (SV-MRS) can be used to differentiate the lipid components. However, it has a limited volume of interest. Alternatively, conventional magnetic resonance spectroscopy imaging (MRSI) methods can be used to cover an entire cross-section over the region of interest, but it requires a long scan time between ~17 to 51 minutes.^30–34^ Acceleration of MRSI sequences had been achieved by reducing the flip angle and repetition time (TR),^35^ or by using a gradient-echo sequence with several echo time (TE) step increments.^36^ These accelerated methods allow a very high resolution (<0.1 mL) within a shorter acquisition time (~10-15 minutes). However, they compromise on large water residual and only allowing for detection of the most intense methylene peaks.^37^ MRSI acceleration was also achieved by using a circular sampling acquisition,^38^ or by implementing an echo planner acquisition technique to speed up the scan by trading the signal-to-noise ratio.^37,39^ However, these techniques do not provide water information without a separate extra measurement.

Providing both, water information with a high spatial resolution is important. Instead of using an external quantification reference or an internal reference located in a different voxel such as bone marrow, which needs B_1_ correction, the water peak information can be used as an internal reference to scale the data.^40^ Additionally, since water has a high signal-to-noise ratio, it is used to determine the local magnetic field shifts and to correct the spectral phase and eddy current effects.^41,42^

Spatial resolution is also important since it enhances the spectral line separation and eventually the detectability of different lipid peaks. The musculoskeletal lipid exists either outside the muscle cells and thus is called extramyocellular lipid (EMCL), or exists as droplets inside the muscle cells and is called intramyocellular lipid (IMCL). EMCL and IMCL are usually separated by ~0.2 ppm due to the local bulk magnetic susceptibility (BMS).^20,21^ Unlike IMCL, EMCL extends along the muscle fibers, and thus its chemical shift may be affected by the fiber orientation relative to the main magnetic field (B0) direction because of the experienced anisotropic BMS.^21,43^ The separation of EMCL from IMCL was found to be maximal when the fiber orientation is parallel to B0. However, the precision frequency of the EMCL starts approaching that of IMCL as the muscle fibers orientation deviates away from B0 direction.^21^ Therefore, muscles with asymmetrical fibers orientation distribution such as the soleus muscle will have broader EMCL spectral linewidth.^44^ Thus, using techniques of high spatial resolution is desired to reduce the potential variation of the fiber orientation within the same voxel and eventually mitigate its influence of widening the EMCL spectral line width. This is especially useful to resolve the peak of extramyocellular L15 and its adjacent smaller peak of intramyocellular L13.

Therefore, there is a need for a reliable and fast non-invasive in vivo quantification method that is capable of providing the spatial distribution for each lipid of interest within a short scan time. In this work, we demonstrate a high-resolution, density-weighted concentric ring trajectory (DW-CRT) metabolite cycling (MC) MRSI acquisition technique to provide the needed fat quantification technique. The DW-CRT k-space-filling technique^45^ allows results of higher resolution (0.25 mL) within a shorter scan time. In addition, by implementing the MC acquisition technique,^46^ simultaneous out phased upfield and downfield (relative to the water frequency) spectra can be provided from a single acquisition. These spectra can be then used to reconstruct separate metabolites and water only spectra. By using this advantage, the water signal information can be used as an internal reference to calculate the FF voxel-wise in a similar approach used by the Dixon method, but for each lipid component individually based on their amplitude and unique resonance frequency. A second objective is to investigate the regional distribution of each lipid component over the calf muscles in an adolescent population.

## 2 METHODS

### 2.1 Human subjects

In vivo calf muscle scans were performed for five healthy non-obese adolescent volunteers [2 males and 3 females; age 12-16 years (median = 14 years); body mass index (BMI) = 20 ± 3 Kg/m^2^]. The scans were acquired at the maximum circumference of the lower leg. All subjects stated that they did not exercise for at least 24 hours before their scan. The study was conducted in accordance with the institutional review board of Purdue University. Before being scanned, an informed written assent was obtained from all the subjects, and written consent was obtained from their parents.

### 2.2 MR scanning parameters

The data were acquired by using the integrated body coil of the Siemens Prisma 3T MR system (Siemens, Germany). The FID DW-CRT MRSI was prescribed using a Hanning-window and the following parameters: field of view (FOV) = 240 × 240 mm^2^, matrix size = 48 × 48, acquisition delay = 4 ms, repetition time (TR) = 1 s, points-per-ring = 64, temporal samples = 512, resolution = 5 × 5 × 10 mm^3^ (0.25 mL), number of rings = 24, spatial interleaves = 4, and spectral bandwidth = 1250 Hz. For the MC, similar parameters to that used in Steel et al. were implemented: an 80 Hz transition bandwidth (−0.95 < M_z_/M_0_ < 0.95) and 820 Hz inversion bandwidth (−1 < M_z_/M_0_ < −0.95), 70 to −750 Hz) downfield/upfield from the carrier frequency (carrier frequency offset = +60 Hz and −60 Hz for downfield and upfield) were defined.^45^ The number of averages was 1, corresponding to a total acquisition duration of 3 minutes and 16 seconds.

For comparison, imaging with a 3-point fast-spin-echo Dixon sequence was performed with echo time (TE) = 11 ms, TR = 5 s, 2 averages, FOV = 200 × 200 mm^2^, and resolution = 0.6 × 0.6 × 10 mm^3^. Since Dixon is considered the standard MR technique to quantify FF and map the fat only and water only distributions, its results will serve as a reference to assess the goodness of the MRSI results.

To get an anatomical image suitable for segmentation, an image was acquired with a T1-FLASH sequence with TR/TE = 250 ms/2.46 ms, flip angle = 60°, 2 averages, 0.6 × 0.6 × 10 mm^3^ resolution, and FOV = 200 × 200 mm^2^.

All three sequences were planned to collect data from the same axial slice placed at the scanner isocenter.

### 2.3 MRSI post-processing

The MRSI data were reconstructed in MATLAB (MathWorks, Natick, MA, USA). The gridding and the Fast Fourier Transform were performed by utilizing the Nonuniform FFT (NUFFT) method^47^ without using any post-hoc density compensation since DW-CRT is already weighted by design.^48^ The voxel-wise frequency and phase corrections were performed using cross-correlation and least-square fit algorithms, respectively, as described in Emir, et al..^49^ The FIDs were smoothed by using a Gaussian filter of 250 ms timing parameter and zero filling to 1024 time points. Following, the water-only and the metabolite-only spectra were created by summing and subtracting the alternating FIDs, respectively, as described in Figure 1.

**FIGURE 1.**
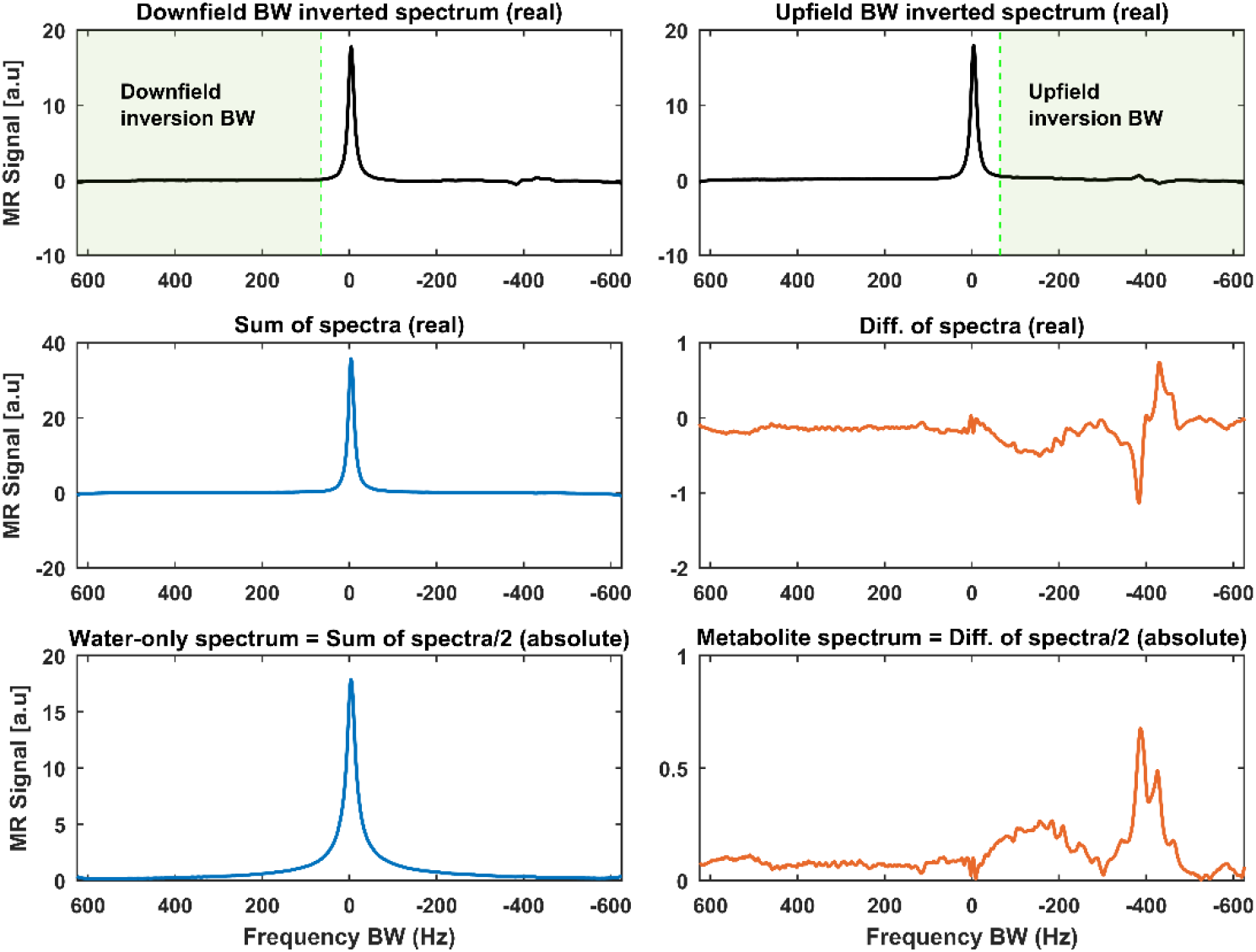
Example of how to get water-only and metabolite-only (includes the lipid peaks) spectra using metabolite cycling (MC) acquisition. The MC acquisition technique includes two selective adiabatic inversion RF pulses, each with a transition over the water bandwidth (BW). The first adiabatic pulse inverts the downfield BW relative to the water frequency (top panel, left), while the second one inverts only the metabolites upfield of the water frequency (top panel, right). The sum of these spectra provides a pure water spectrum with a minimal residual metabolite signal (middle panel, left), while their difference gives a pure metabolite spectrum with insignificant residual water (middle panel, right). The final spectra are magnitude spectra and divided by two since the summation and subtraction give double the original signal (bottom panel)

### 2.4 Quantification

The Dixon technique provides a water image and a total lipid image. In order to generate a FF map out of these Dixon images, the following formula is used:

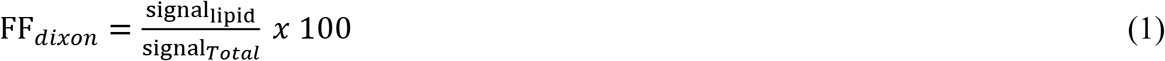

Where signal_*Total*_ = lipid + water images signals.

As for MRSI, it provides water-only and metabolite-only resonance spectra. Following their spectral post-processing, these spectra from each voxel are passed into LCModel to fit each peak of the spectra individually and return their integrated signal.^50^

In order to avoid phase correction artifacts in areas overwhelmed by lipids such as bone marrow or subcutaneous fat regions, the LCModel phase correction option was set to zero. Instead, the magnitude value was used as previously done by Meisamy et al..^51^ In order to correct for the long water T_1_ relaxation time effect, the water reference attenuation correction parameter, ATTH2O, in LCModel was used. This parameter was determined based on the T_1_ signal relaxation term, 1-exp(-TR/T_1_), where the T_1_ value of water in skeletal muscle was assigned to 1412 ms as measured by Stanise et al. at 3T.^52^ LCModel’s basis set of muscle spectra, “muscle-5”, was used to fit the magnitude MRSI-spectra. To construct the MRSI maps, only peaks with Cram’er-Rao lower bounds (CRLB) values of 8% or less (measured by LCModel) were used. The FF was then calculated by using a similar expression to that used to calculate the FF_*dixon*_:

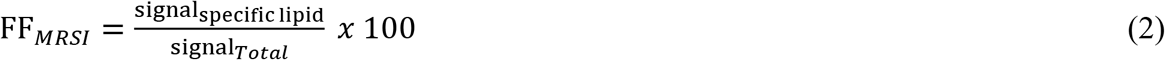

For both techniques the water fraction (WF) is calculated by the following formula:

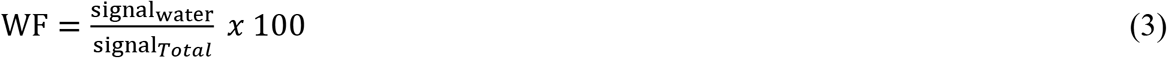

In addition, to assess the MC performance within the calf area, the residual water (RW) fraction was calculated as:

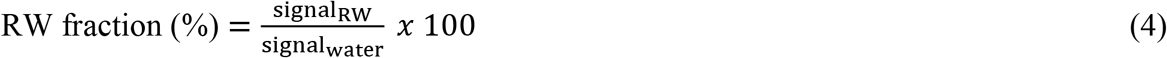

Where signal_RW_ is the signal of the residual water peak within the metabolite-only spectra fitted by the LCModel.

### 2.5 Muscle segmentation

To assess the spatial FF distribution of each lipid component within the calf muscles, muscles were manually segmented by drawing ROIs over each of the eight main muscles based on their high-resolution T1-weighted image. The ROIs borders were determined by following the muscle boundaries. These ROIs were down-sampled to match MRSI resolution and co-registered to each lipid FF map to assess their distribution voxel-wisely. To avoid any partial volume effect by adjacent structures, voxels on the borders were excluded out of the down-sampled ROIs (Figure 2).

**FIGURE 2.**
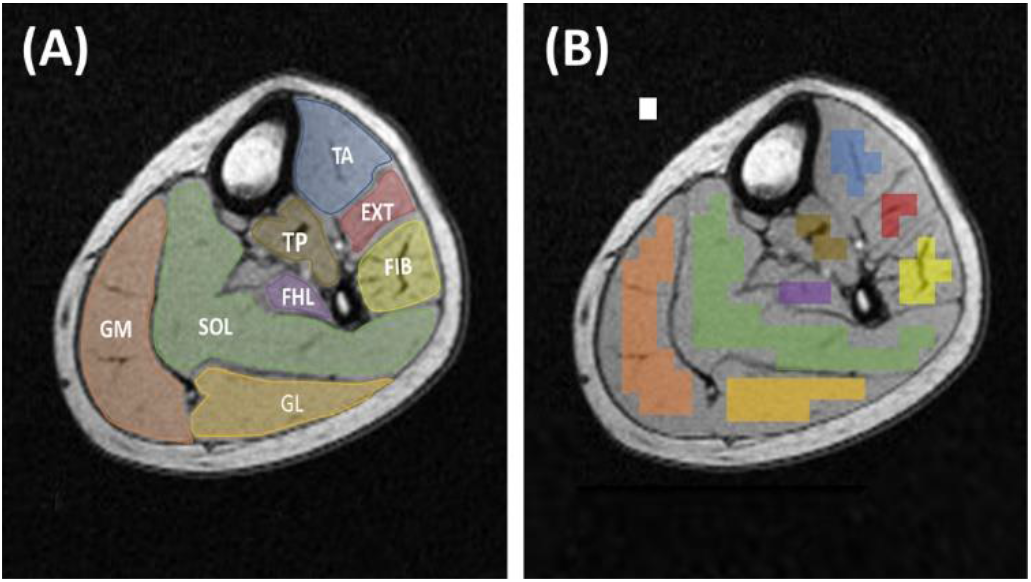
A, Calf Muscle region of interests (ROIs) drawn on a high-resolution T1 axial image, which provides a good anatomical delineation. Eight ROIs were used to cover the main eight calf muscles. SOL, Soleus muscle; FIB, Fibularis muscles; EXT, Extensor longus muscles; TA, Tibialis anterior muscle; GM, Gastrocnemius medialis muscle; GL, Gastrocnemius lateralis muscle; FHL, Flexor halluces longus muscle; TP, Tibialis posterior muscle. B, The same ROI set after being down-sampled into MRSI resolution and removing the voxels on the muscle borders. The white box represents one ROI voxel

### 2.6 Data analysis

To compare the results of the two used techniques, the Structural Similarity (SSIM) Index method,^53^ implemented in MATLAB, was used to find the level of similarity in fat detectability between the FF maps generated from the MRSI versus those from the Dixon technique. SSIM analysis compares two images and returns their signal intensity and structural similarity level (index) for each voxel as well as a global (mean of all voxels) value.

Additionally, in order to assess each lipid component FF spatial distribution within the muscles estimated by the proposed MRSI method, the previously segmented muscle ROIs were used with each lipid component FF map. To eliminate outlier voxels from generating false-positive findings, a minimum positive voxel threshold (MPVT = 1/number of subjects = 20% of the ROI voxels from all subjects) was considered. This means that only muscles with enough (i.e., above MPVT) positive voxels (voxels containing the lipid of comparison and passed the CRLB criteria of peak fitting goodness) were included in the regional comparison. To conduct this regional comparison, the Kruskal Wallis one-way analysis of variance test was used to assess the lipid spatial distribution variation and flowed by the Bonferroni multiple comparison test to determine whether a significant FF difference exists between any two muscles.

## 3 RESULTS

As shown in Figure 3, generated water- and lipid-only maps of MRSI are in agreement with the anatomical distribution of those provided by the Dixon technique. The proposed MRSI technique further provided two sets of spectra from one acquisition, with the first set being pure water spectra, and the second set containing spectra with information about different lipid components and other metabolites. The generated MRSI spectra were fitted by LCModel, which identified eight lipid peaks found within the metabolite spectra (L09, L11, L13, L15, L21, L23, L53, and L55) plus Cr30, Cr39, and Crn32 (see examples in Figure 4). As can be seen in Figure 5, eight separate FF maps of each lipid component were reconstructed from the quantification results by LCModel for the fitted lipid and water peaks. The MC method resulted in only 1.3 ±1.2% RW fraction that was only found in half of the calf voxels (Figure 6).

**FIGURE 3.**
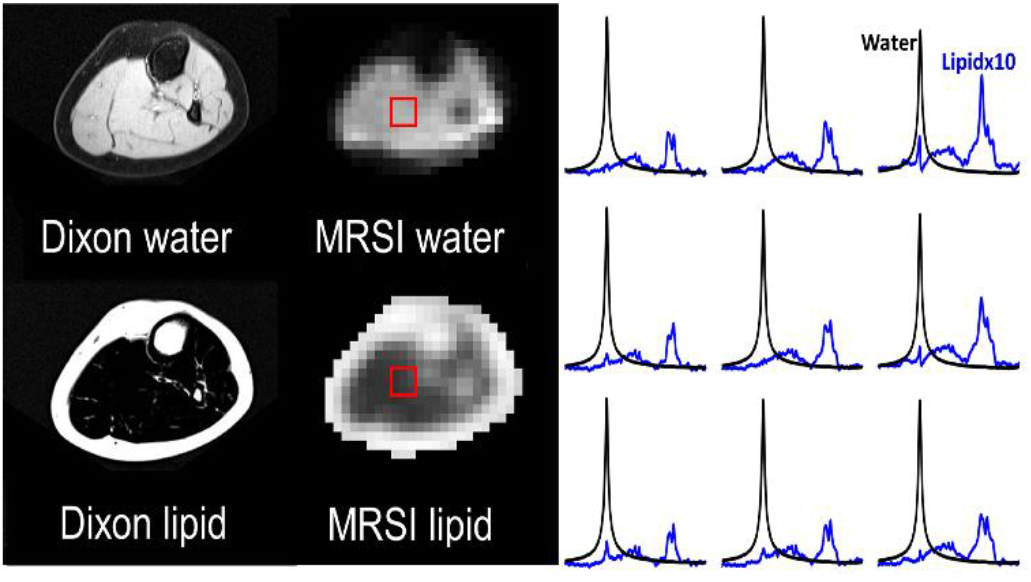
Dixon versus MRSI output. The results of the FID density-weighted concentric ring trajectory (DW-CRT) metabolite cycling MRSI sequence are in line with the Dixon images. On the right, representative water (black) and lipid (blue) spectra acquired from the same location (box) by MRSI. The lipid peaks were magnified ten times compared to the water peak for better visualization

**FIGURE 4.**
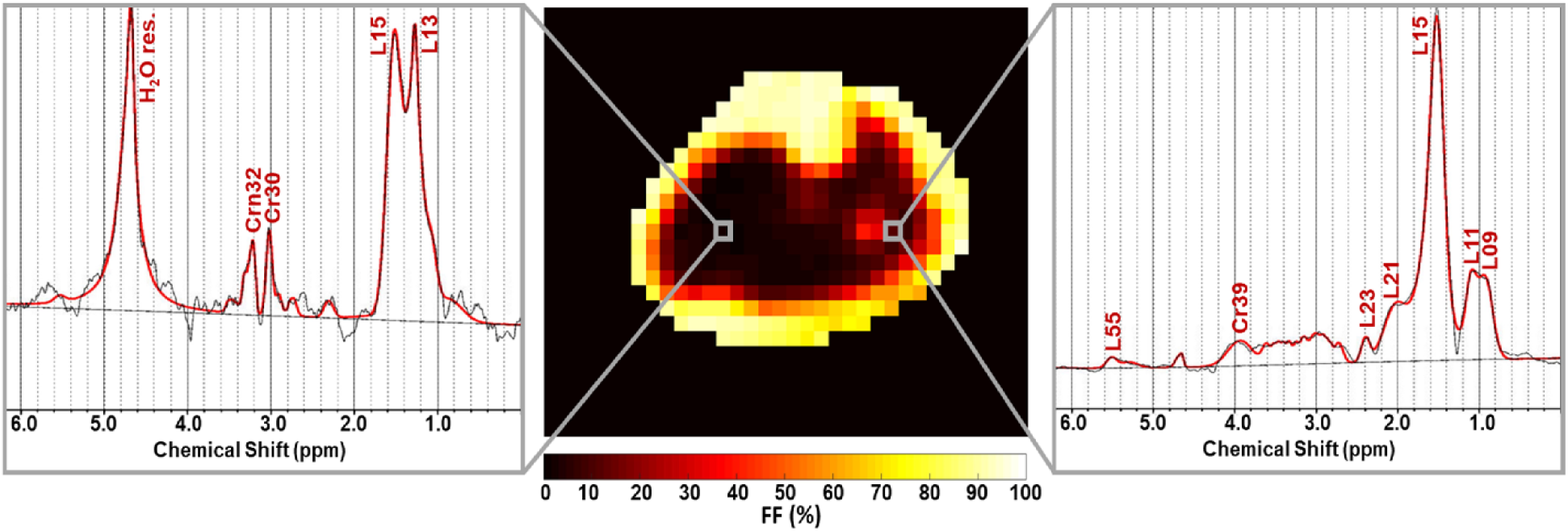
Representative spectra fitted by LCModel. The shown spectra (black) were acquired from the voxels highlighted on the total fat fraction map. The LCModel fit is shown in red, with the fitted lipids and metabolites labeled. H_2_O res. stands for the residual water signal. Other metabolites than lipids can also be detected such as CH_3_ and CH_2_) groups of creatine that resonate at 3.0 ppm (Cr30) and 3.9 ppm (Cr39), respectively, in addition to the CH3 group of carnitine (Crn32)

**FIGURE 5.**
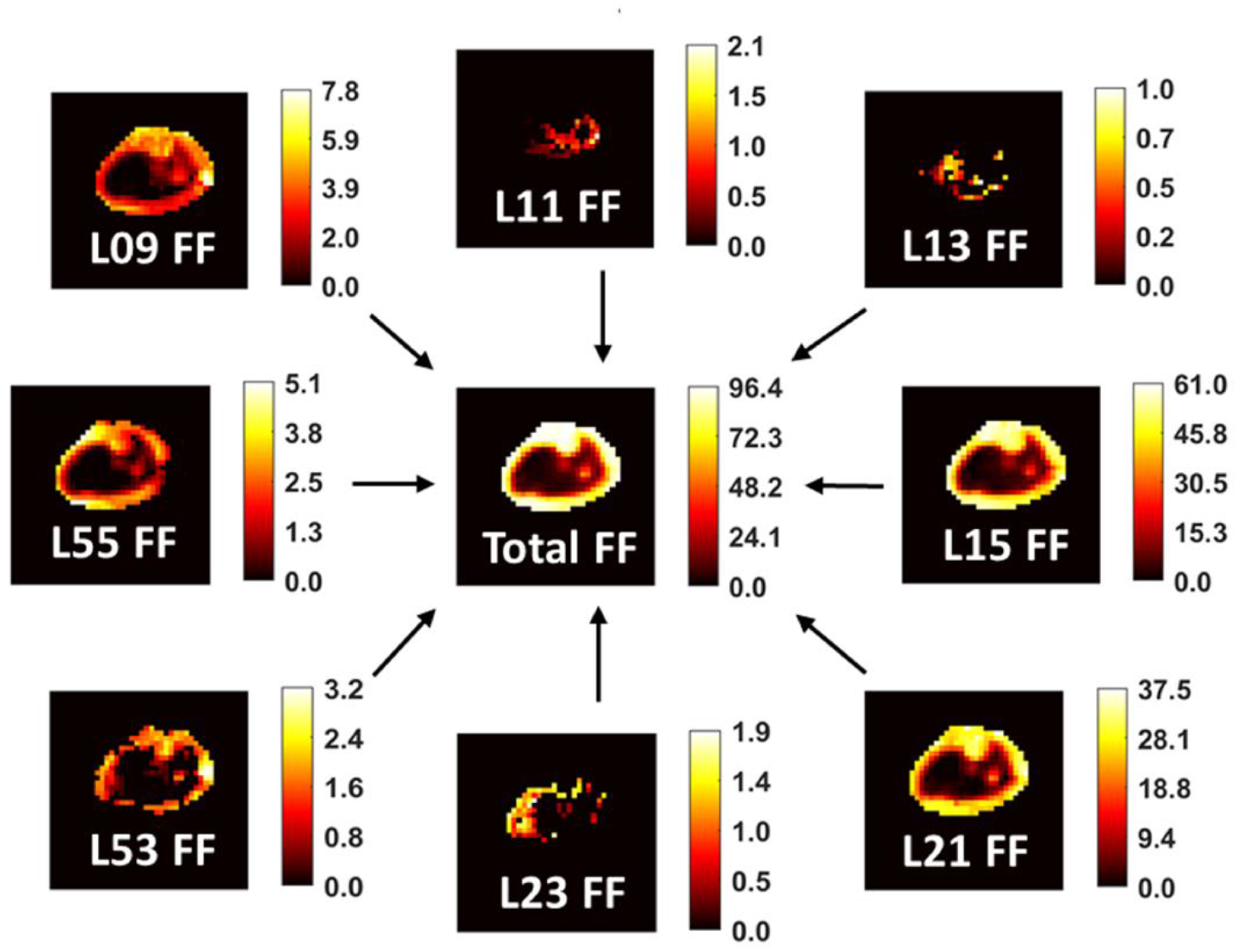
Representative fat fraction (FF) maps for each lipid component that was fitted by LCModel. Only results with CRLB of 8% or less were included. The scale next to each map indicates the FF values from 0 to maximum in percent

**FIGURE 6.**
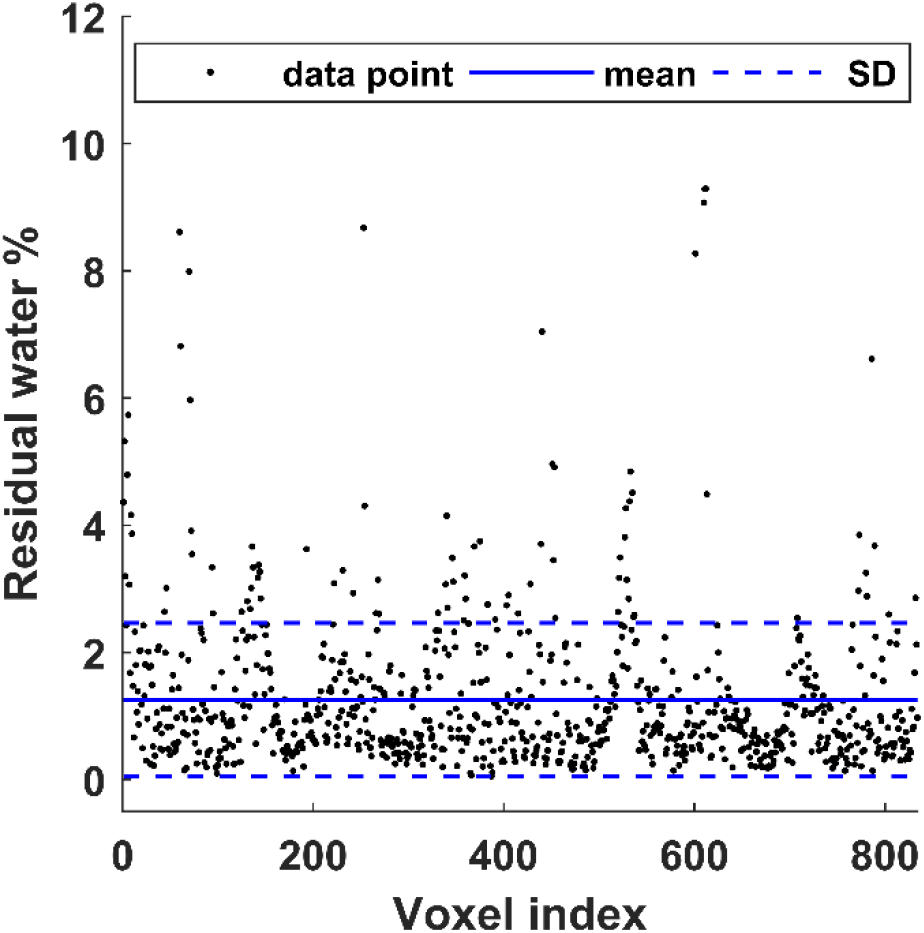
Residual water (RW) after implementing the MC. Out of 1626 voxels within the calf area (from the five subjects), the LCModel could fit residual water in 832 voxels with a mean RW fraction of 1.3 ± 1.2%

The results of the SSIM analysis comparing the FF_*dixon*_-map to each lipid component FF-map generated by MRSI are illustrated in Figure 7. The data presented as SSIM index mean ± standard error that was evaluated for all the five scanned subjects. Figure 8 demonstrates an example of the SSIM analysis result from one subject’s data. The example involved the lipid of the highest signal, the L15, and the lipid of the lowest signal, the L13. In this example, the mean SSIM index results from comparing the Dixon technique to the L13 results was very low (0.03) compared to the similarity of L15 (0.60). In fact, the Dixon technique could not detect L13 and L15 in several locations where the MRSI was able to detect considerable signals from these lipids. For demonstration, in the same figure, spectra from 2×2 voxels covering a random area where total mismatching present (SSIM = 0) were shown.

**FIGURE 7.**
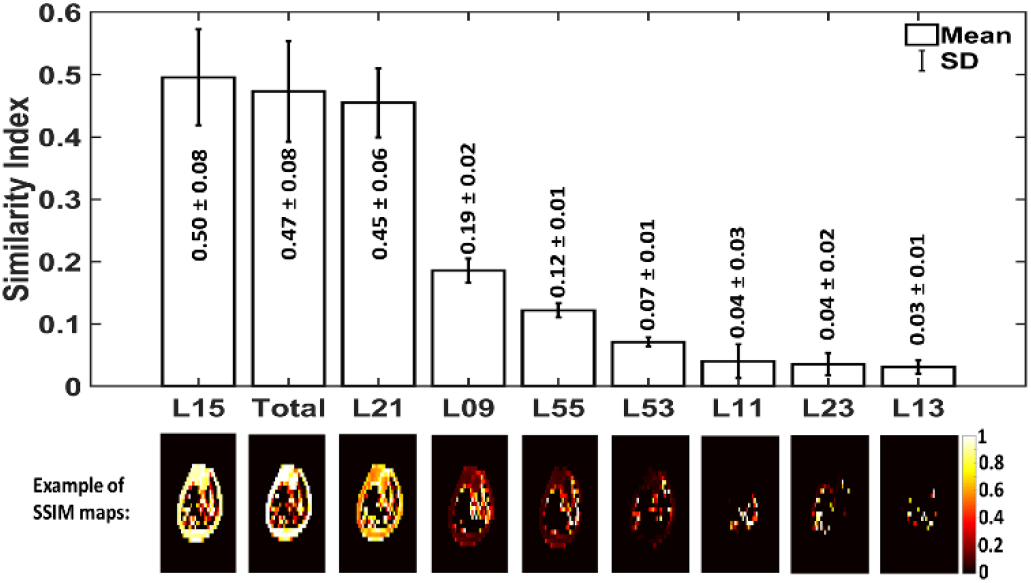
Dixon similarity to MRSI maps results. The structural similarity (SSIM) indices mean as a result of comparing the Dixon fat fraction (FF) map to the FF-map of each lipid detected by MRSI. The mean and standard error were calculated based on the data from the five healthy subjects. The SSIM indices range between 0 and 1. SSIM=1 represents a perfect similarity. The results are ordered based on their order of similarity from the highest to lowest (left to right). Representative SSIM maps from one subject are depicted below each bar of their corresponding lipid component. The results suggest that Dixon’s fat signal is mainly coming from L15, L21, and L09

**FIGURE 8.**
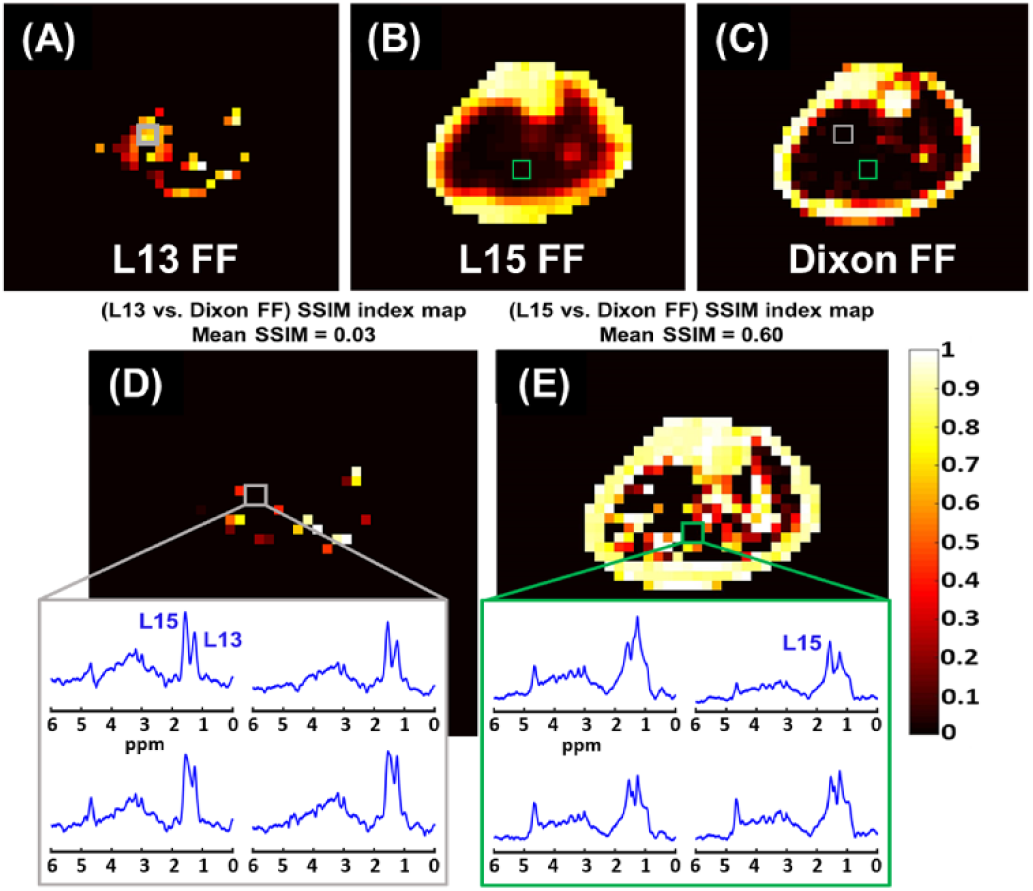
Fat fraction (FF) distribution maps of MRSI L13 (A), MRSI L15 (B), and the Dixon-MRI undifferentiated-fat-fraction image (C). The Dixon MRI image was down-sampled to MRSI resolution for a better comparison. The corresponding SSIM index maps show the structural similarity (SSIM) between the Dixon FF image and the MRSI L13 FF map (D), and the L15 FF map (E). The mean SSIM value is listed above each SSIM map. The dark areas of SSIM = 0 on the maps (box) represent total mismatching between MRSI and Dixon results. Within these areas, only MRSI could identify lipid peaks

As shown in Figure 9, L09 FF was present in all the muscles that were defined in Figure 2 except in the GL as L09 was detected within less than 20% of its ROI voxels. According to the Kruskal-Wallis (KW) statistics, the FF of L09 was highly variable among the tested muscles (KW-P < 0.001), and it was significantly lower in GM than in EXT (*P* < 0.001), FBL and SOL (*P* < 0.01) based on the multi comparison statistics. For L11, it existed within less than 20% of the GM voxels. The KW statistics also indicated a very high variation of this lipid among the other muscles (KW-*P* < 0.001). The L11 level was significantly lower in TA compared to FHL (*P* < 0.001), TP (*P* < 0.01), SOL and FIB (*P* < 0.05), and it was also lower in GL relative to FHL (*P* < 0.01). For L13, the MPVT was exceeded only in five muscles (EXT, FHL, GL, GM, and SOL). Its level among these muscles was only higher in SOL compared to it within GM (*P* < 0.001). On the other hand, L15 was detected within all the calf muscles with a very large level variation (KW-*P* < 0.001). The multi-comparison analysis detected that L15 has a higher FF in EXT and FIB compared to GL, GM, and SOL (*P* < 0.001). The L15 FF was also smaller in FHL compared to FIB (*P* < 0.001) and EXT (P < 0.01) and smaller in GM compared to TP (*P* < 0.01) and TA (*P* < 0.05). Similar to L15, the L21 was also detected within all the muscles with a strong FF distribution variability (KW-*P* < 0.001), with a significantly higher FF within FIB than in GM (*P* < 0.01), and SOL (*P* < 0.05), and EXT has higher FF compared to GM and SOL as well (P < 0.05). On the other hand, L23 was detected with enough voxels larger than the MPVT only in the three muscles located at the posterior part of the leg (GL, GM, and SOL) and without significant FF variation among them (KW-*P* = 0.2). For L53, The MPVT was satisfied in only five muscles (EXT, FIB, FHL, GM, and TA) without significant fat level variation (KW-*P* = 0.4). The L55 was not above the MPVT for the FHL and GL voxels. Among the other muscles, the L55 FF distribution was found to be moderately variable (KW-*P* = 0.02) as only SOL has significantly higher FF of L55 relative to TP (*P* < 0.05).

**FIGURE 9.**
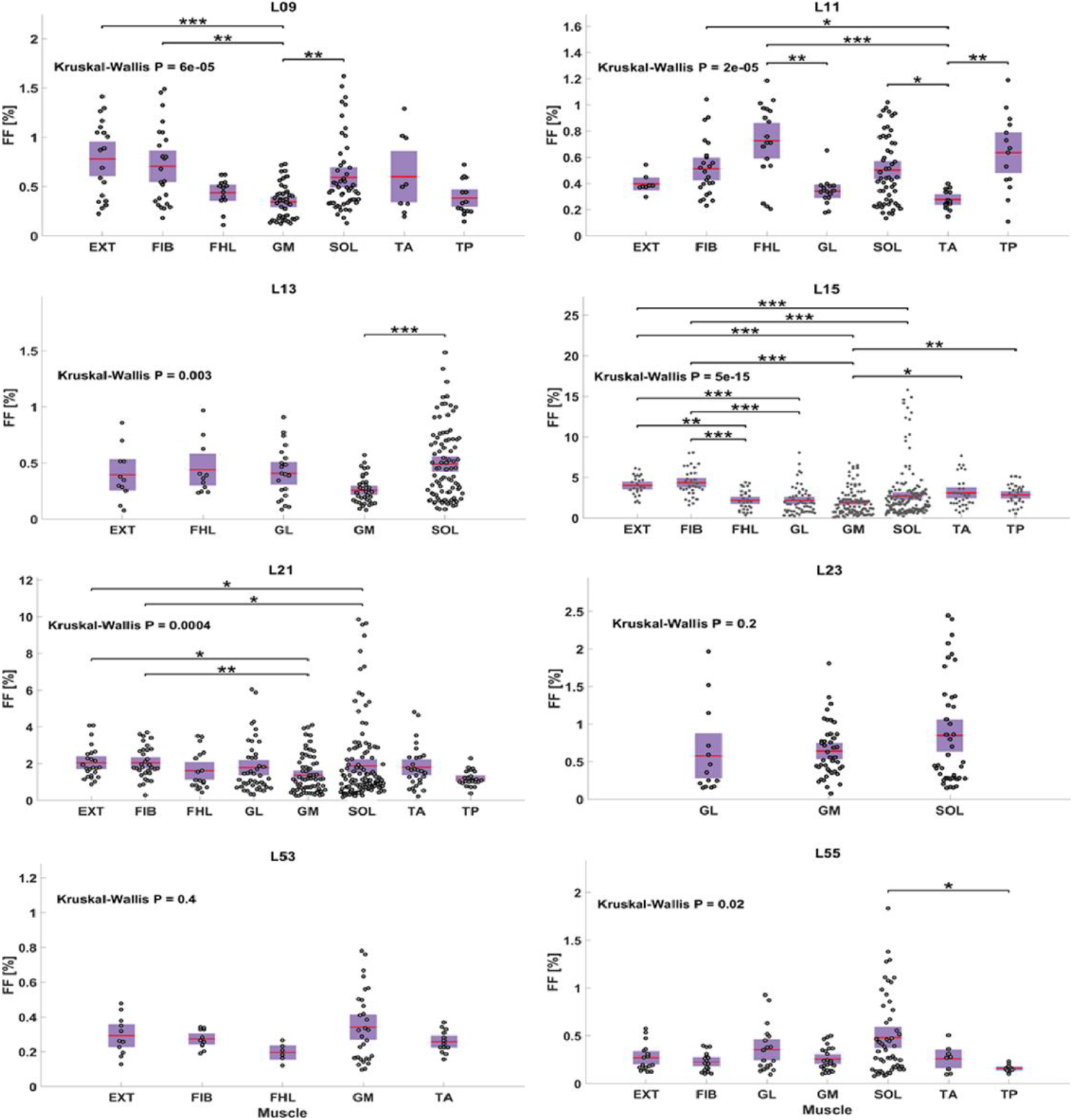
Regional comparison of each lipid component fat fraction (FF) distribution within the calf muscles. Each plotted point represents one voxel FF data. Only muscles with at least 20% of their voxels containing the lipid of comparison are included. The p-value (P) of Kruskal Wallis analysis of variance test is listed for each lipid distribution. The *, **, *** represent *P* <0.05, *P* <0.01, and *P* <0.001 respectively, and are shown when a significant difference exists between any two muscles according to the Bonferroni multi-comparison test

## 4 DISCUSSION

Our proposed technique has a set of advantages. It is fast, which reduces motion artifact, and reduces the MRI scan expenses. It has a high spatial resolution to reduce the degrading influence of the point spread function and the fiber orientation heterogeneity on spectral quality. Furthermore, it has a short TE relative to the lipid and water T2 values, so quantification is possible without the need for a T2 correction. Most importantly, it provides water-only and separate lipid signals for more precise FF quantification that eventually reflected in better diagnosing ability. The water information is provided simultaneously to be used as an internal reference and to perform post-processing corrections without the need for an extra acquisition by using the MC.

Additionally, this work has shown that clinically important lipids, but of relatively low MR signal such as L13 are better evaluated by the proposed MRSI method rather than the imaging technique. This recommendation is based on the findings observed from the SSIM analysis. As demonstrated in the results, the MRSI method showed a better performance identifying those lipids of lower levels. Such molecules of interest normally have a low signal that is hard to be detected by imaging techniques or overwhelmed by larger signals from other molecules within the same voxel. Indeed, the lipid signal measured by the conventional Dixon techniques mainly originates from the methyl and methylene peaks. This finding agrees with a previous phantom study that showed that the Dixon method provides lower accuracy in quantifying total FF compared with MRS when the signal from all lipid peaks was summed.^54^

We used a conventional Dixon technique that is similar to the most common Dixon sequences available with the most clinical scanner. Since we used the body coil, we had to turn off the parallel imaging feature, which resulted in prolonging the scan time. To mitigate the impact of not using the parallel imaging, we used a fast spin-echo sequence^55^ and adjusted its parameters (use short TE and long TR) to make it a proton-density weighted. Although that other advanced Dixon techniques were introduced and some of them suggested methods to account for the fat multipeak existence such as the work done by Yu et al.,^56^ they are still not able to allocate different map for each lipid of interest like the proposed MRSI technique. Indeed, some studies comparing Dixon techniques to SV-MRS, suggested MRSI as a promising fat quantification technique as the SV-MRS showed higher sensitivity and dynamic range compared to Dixon, but it has lower resolution and spatial coverage.^57^

We decided to perform this study on an adolescent population to investigate the lipid distribution in this age group as several studies have conducted a similar assessment but on adult populations.^30,58,59^ Most of these adult studies considered only L13 and L15 distribution. Our findings agree with all these previously published studies for L13. However, forL15, our findings on adolescents agree only with one study findings^58^ that there is no significant variation in L15 content within SOL and TA. For the other three studies, the L15 level was always significantly larger in SOL relative to it in TA. Although that our L13 distribution conclusion matches the published studies, the other studies were able to report values for L13 level in TA, which usually the lowest among the investigated muscles, but our findings suggest that no L13 was detected in this muscle. Potential reasons for these contrast in findings could be the age impact as muscles lipid content found to be lower in younger age,^60,61^ because some of these studies conducted by using a large SV-MRS, which is more sensitive to the partial volume effect, or MRSI without considering a method to exclude the false positives. In addition, SV-MRS was found to have more variability to reproduce the same measurement compared to MRSI based on the study done by Shen et al.^62^ on the TA muscle which can be another potential reason for this difference in findings of this muscle. The uneven distribution of L13 across muscles was related to the amount of slow-twitch (type I) fibers in the muscles.^30,58^ For instance, the majority of SOL fibers are of type I, whereas GM and GL contain more fast-twitch (type II) fibers.^63^ In contrast to type II fibers, type I is smaller in diameter, contains more mitochondria, depends less on the ATPase activity to produce energy and thus stores more IMCL for the mitochondrial oxidation process.^63–65^ In addition, the functional role^58^ and fiber orientation^31^ of muscles were suggested as a potential contributor in the lipid content regional differences. Thus, the reason for the contrast of the lipids level among muscles still needs further investigations.

Conducting this research by scanning healthy young subjects was challenging since their tissue fat content is not as high as that in obese people or patients with a condition associated with a fat level increase. Further, using the body coil boosted the challenge level since it has lower detecting sensitivity, which is very important for spectroscopy applications. However, our technique could successfully produce high-quality data. Even so, the results can be enhanced further by using another smaller coil with multi-channels instead of the body coil. Indeed, by using the scanner-integrated body coil, we showed the feasibility to use it to achieve good spectra while another coil can be attached to the scanner for any other purpose that is not a rare situation. For instance, it can be used with another non-proton coil to investigate other nuclei such as phosphorus, carbon, or sodium without the need to reposition the subject. Furthermore, the data quality can be increased more by using a shorter TE that helps to reduce the eddy current effect on the smoothness of the spectra baseline.

A summary of the latest MRI/S methods for neuromuscular disease monitoring has been recently published.^66^ Within this publication, the importance of advancing fat quantification techniques was emphasized. We think that our proposed technique would help the field by providing a different approach to assess fat infiltration diseases. Actually, the importance of this proposed technique can be extended beyond the focus of this work to cover more applications such as quantifying FF in the bone marrow to assess osteoporosis,^29^ and Anorexia nervosa,^67^ or to assess the intrahepatic lipid (IHL) in the liver, which is used to evaluate the hepatic steatosis.^68^ Nonetheless, our proposed technique can be used to assess physiological change by mapping extra metabolites such as Crn32 that associated with muscle exercises, or maybe implemented to assess choline level to evaluate hepatic and breast tumors.^69–71^ Further, this MRSI sequence could fill the need for a fast fat quantification for fatty liver since it located in a moving location due to breathing. This need was expressed in Parente et al. study, where they preferred imaging technique since SV-MRS required a long time, and it may increase the chance of misinterpretation in cases of heterogeneous steatosis.^72^

To the best of our knowledge, this is the first time a technique provided to generate separate spatial FF maps for the lipid components. However, the lower spatial resolution of MRSI relative to imaging may be a limitation for some applications. Thus, a technique to improve the MRSI spatial resolution further would be useful to serve any fat quantification application.

## CONCLUSIONS

The proposed MRSI technique provides a needed tool to quantify different lipid components non-invasively. It can reliably detect fat components with high sensitivity and provides pure water information to calculate the FF accurately. Most importantly, it can allocate separate quantitative spatial maps for the lipid components over the entire region of interest within clinically feasible acquisition time, ~ 3 minutes.

## ACKNOWLEDGMENTS

The study was supported by the Indiana CTSI and funded in part by grant #UL1TR001108 from the NIH, NCATS, CTS Award, as well as a pilot grant by the College of Health and Human Sciences, Purdue University.

